# Charting sex divergence in the human brain across the lifespan

**DOI:** 10.64898/2026.10.11.758308

**Authors:** Minhui Zhu, Lianglong Sun, Mingrui Xia, Yanchao Bi, Qingjiu Cao, Rui Chen, Yuan Chen, Taolin Chen, Yuqi Cheng, Zhengjia Dai, Yao Deng, Yuyin Ding, Qi Dong, Jia-Hong Gao, Qiyong Gong, Ying Han, Zaizhu Han, Chu-Chung Huang, Hao Huang, Ruiwang Huang, Ran Huo, Lingjiang Li, Qixiang Lin, Bangshan Liu, Chao Liu, Hesheng Liu, Ningyu Liu, Ying Liu, Yong Liu, Jing Lu, Leilei Ma, Weiwei Men, Chenxuan Pang, Shaozheng Qin, Wen Qin, Jiang Qiu, Shijun Qiu, Tianmei Si, Shuping Tan, Yanqing Tang, Sha Tao, Dawei Wang, Fei Wang, Jiali Wang, Pan Wang, Qian Wang, Xiaoqin Wang, Yanpei Wang, Dongtao Wei, Yankun Wu, Peng Xie, Xiufeng Xu, Li Yang, Chunshui Yu, Huishu Yuan, Haibo Zhang, Xi Zhang, Gai Zhao, Kun Zhao, Yanting Zheng, Alzheimer’s Disease Neuroimaging Initiative, MCADI, DIDA-MDD Working Group, Xinyuan Liang, Yong He

## Abstract

Biological sex fundamentally shapes human brain organization, but sex-specific normative neuroanatomical trajectories across the lifespan remain largely uncharted. Here, we constructed independent, sex-specific lifespan brain charts using structural neuroimaging data from 59,915 healthy individuals (29,760 males and 30,155 females) ranging in age from 266 postconception days to 100 years. By examining 296 structural phenotypes across global, cortical, and subcortical measures, these models revealed widespread sex differences in maturational timing, with males reaching peak milestones later than females. These trajectories demonstrate that sex differences evolve dynamically, with phenotype-specific windows of emergence and maximal separation. Compared with conventional sex-pooled references, sex-specific models achieved superior predictive accuracy and reduced misestimation of individual deviations in healthy populations. Across five neuropsychiatric conditions, sex-specific models improved the detection of extreme deviations and revealed both shared and sex-dependent patterns of disorder-related neuroanatomical abnormalities. These sex-specific charts establish tailored normative references for assessing brain development, ageing, and disease.

## Introduction

Biological sex fundamentally shapes the structural and functional organization of the human brain ^1–6^, driven by a lifelong interplay of sex chromosomes ^7, 8^, gonadal hormones ^8, 9^, and genotype-by-sex interactions ^8, 10–12^. Delineating how male and female brain trajectories diverge is essential for understanding individual variations in cognition and emotions. Crucially, it also provides a neurobiological foundation for the striking sex biases observed in the prevalence, onset, and pathophysiology of neuropsychiatric conditions ^13^, including autism spectrum disorder (ASD) ^14, 15^, attention-deficit/hyperactivity disorder (ADHD) ^8^, major depressive disorder (MDD) ^16^, and Alzheimer’s disease (AD) ^17^. Despite these profound clinical implications, the spatiotemporal architecture of sex divergence across the lifespan remains largely uncharted.

Over the past two decades, numerous neuroimaging studies have examined sex differences in brain organization. Although males consistently show larger global brain volumes, findings regarding the regional topography and temporal dynamics of sex differences remain highly controversial ^18–22^. Spatially, some studies have reported greater frontal thickness in males ^23^, whereas others have found broadly thicker frontal and parietal cortices in females ^24, 25^, or no significant sex difference at all ^21, 26–28^. Similar discrepancies have been reported for subcortical structures, including the amygdala ^29, 30^, putamen ^25, 29, 31^, and hippocampus ^30, 32^. Temporally, the picture is equally fragmented. Although some studies have suggested that cortical grey matter volume peaks earlier in females ^33, 34^, other studies have reported no such peak differences ^27^ or conflicting results regarding which sex matures earlier in specific areas (e.g., the temporal cortex) ^34, 35^. Even when earlier female maturation is observed, the estimated magnitude of this temporal gap varies substantially ^33, 34^. These discrepancies likely stem from a reliance on restricted age ranges—such as infancy ^36–38^, childhood and adolescence ^34, 39^, or adulthood ^25^—rather than continuous lifespan modelling. This limitation is further compounded by sparse sampling during highly dynamic early developmental stages ^19, 22, 40, 41^ and heterogeneous imaging and analytic protocols ^22^. Notably, even the latest lifespan brain charts built on large-scale imaging datasets ^42–45^ are still constrained in this regard. Most current normative modelling frameworks (e.g., generalized additive models for location, scale and shape, GAMLSS) treat sex simply as a categorical covariate, a practice that forces male and female trajectories to share identical curve shapes and peak ages, thereby precluding any independent estimation of sex-specific milestone timing and population variance. Without fully independent, sex-specific normative models, the true trajectories of lifespan sexual dimorphism remain elusive, ultimately hindering the accurate quantification of neuropathological deviations in clinical populations.

To address these gaps, we constructed fully independent, sex-specific lifespan brain charts using large-scale structural neuroimaging data from 59,915 healthy individuals (29,760 males and 30,155 females), spanning from 266 days postconception to 100 years of age. By applying harmonized processing pipelines and standardized statistical modelling, we delineated sex-specific normative trajectories and milestones for 296 structural phenotypes. We demonstrate that in comparison with conventional sex-pooled models, explicitly modelling sex-specific trajectories not only improves predictive accuracy in healthy populations but also mitigates the misestimation of structural deviations across multiple neuropsychiatric conditions. These sex-specific brain charts establish distinct neuroanatomical timelines and provide a tailored reference framework for the precise, sex-aware assessment of human brain health.

## Results

After rigorous quality control, our final dataset included 59,915 healthy individuals (29,760 males and 30,155 females) from 172 global sites (Fig. 1a; see Methods). To capture sex differences in lifespan brain maturation, we constructed independent normative models for males and females across the 296 structural phenotypes (Fig. 1b). These “sex-specific models” were estimated using GAMLSS, with the scanner modelled as a random effect (Fig. 1c). For comparison, we also generated “sex-pooled models” by fitting GAMLSS to the combined male and female dataset with sex as a categorical covariate, which is consistent with recent brain charting approaches ^42^ (Fig. 1d). We then evaluated both modelling strategies in clinical cohorts comprising 3,914 healthy controls and 5,777 patients with AD, MCI, MDD, ASD, or ADHD (Fig. 1d). This allowed us to compare how accurately each approach characterized typical maturation in healthy participants, and how sensitive it detected disease-related structural deviations in patients. Building on this, we compared case–control deviations for males and females separately, and identified regions that exhibited sex-by-diagnosis interactions (Fig. 1e).

**Figure 1.**
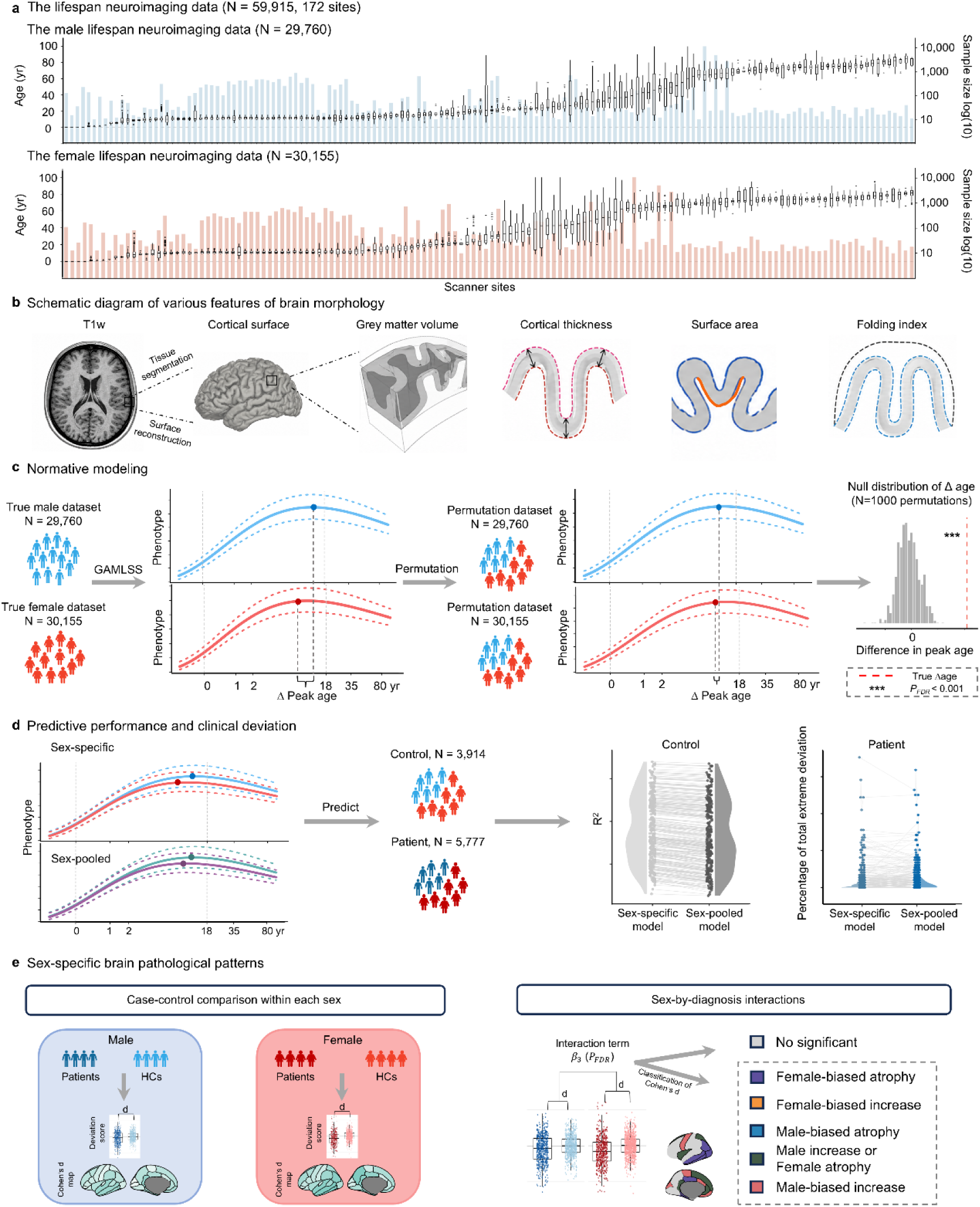
Dataset and workflow of the study. a, Age distribution of male and female participants at each site. **b,** Schematic diagram of brain morphology. **c,** Sex-specific normative modelling. For each brain phenotype, we fitted separate normative models for male and female participants using the GAMLSS framework, followed by permutation testing to evaluate the statistical significance of peak age differences between sexes. **d,** Predictive performance and clinical deviation analysis. We compared sex-specific and sex-pooled models by applying them to out-of-sample data to assess predictive accuracy (R^2^) and to patient cohorts to compare the percentage of extreme deviations. **e,** Sex-specific neuropathological pattern analysis. Sex-specific Cohen’s d maps were derived from case‒control comparisons of the phenotypic deviation scores for males and females. Regions exhibiting significant sex-by-diagnosis interactions were further classified on the basis of the direction of the interaction and the corresponding sex-specific case–control effect sizes. yr, year; GAMLSS, generalized additive models for location, scale and shape; HCs, healthy controls.

### Sex-specific normative growth of global brain phenotypes

While male and female lifespan charts revealed broadly similar, inverted-U shaped trajectories, males exhibited later maturational peaks across most global volumetric and surface-based measures (Fig. 2). Specifically, the total intracranial volume peaked during late adolescence and early adulthood (M: 27.8 years, with 95% confidence interval of [25.4–28.6] years, F: 16.1 [14.9–17.2] years, Δage = 11.7 years; *P_perm_* _<_ 0.001, FDR-corrected; Fig. 2a), whereas the total cerebral and cerebellar volumes peaked in early and late adolescence (cerebrum: M: 12.8 [12.2–13.3] years, F: 10.7 [10.2–11.2] years, Δage = 2.1 years; cerebellum: M: 19.8 [19.2–20.6] years, F: 16.2 [15.5–17.0] years, Δage = 3.6 years; both *P_perm_* _<_ 0.001, FDR-corrected; Fig. 2b, c).

**Figure 2.**
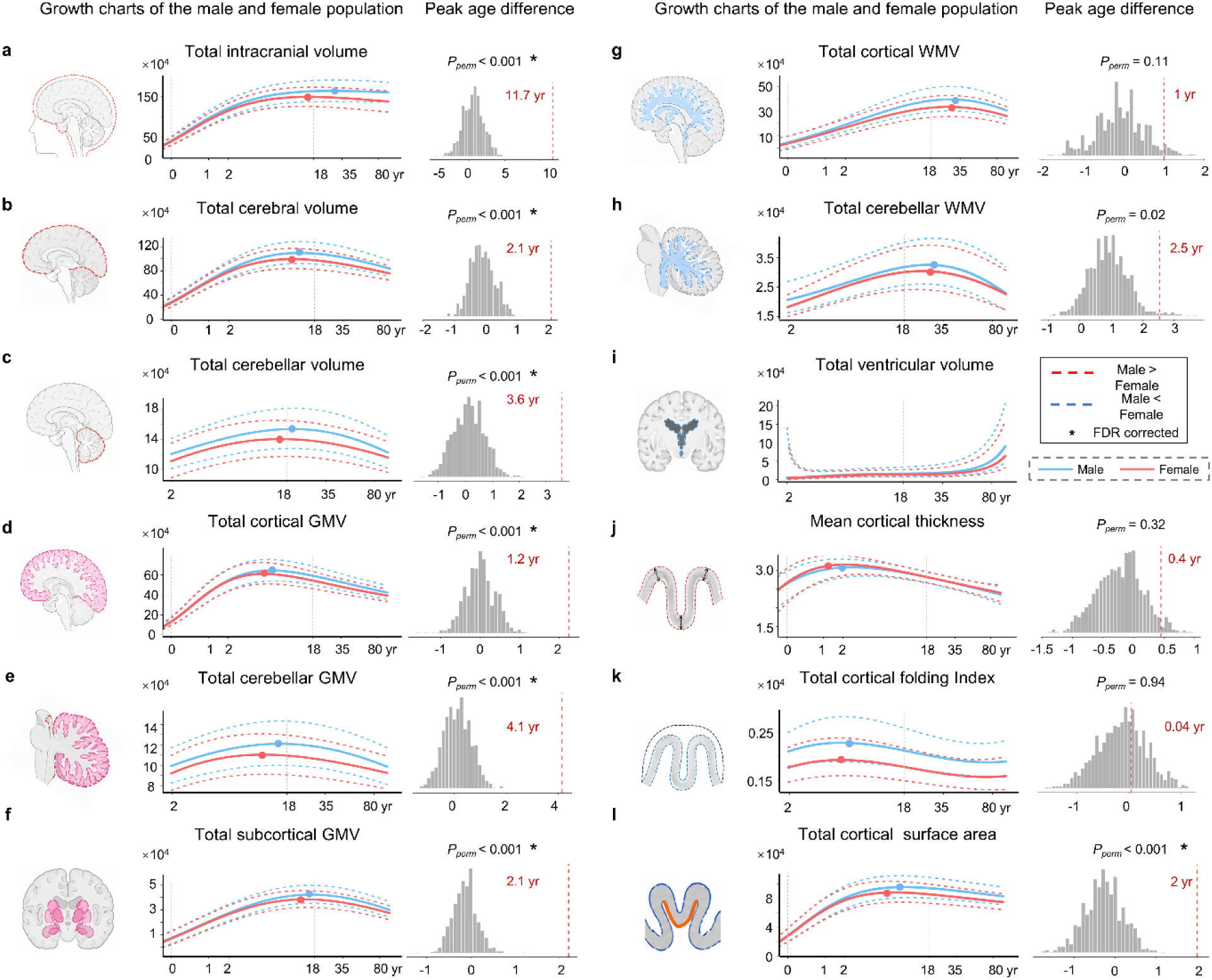
Sex-specific growth patterns of global brain phenotypes. Growth curves of total intracranial volume (**a**), cerebral volume (**b**), cerebellar volume (**c**), cortical GMV (**d**), cerebellar GMV (**e**), subcortical GMV (**f**), cerebral WMV (**g**), cerebellar WMV (**h**), ventricular volume (**i**), mean cortical thickness (**j**), total cortical folding index (**k**), and total cortical surface area (**l**). For each panel, the left side represents the normative growth curves across the lifespan for the males (blue) and females (red) (solid line: median; dotted lines: 5th and 95th percentiles) and the right side represents the null distribution of peak age differences evaluated via a two-sided permutation test (1,000 iterations; see Methods). GMV, grey matter volume; WMV, white matter volume; yr, year.

Grey matter volume (GMV) followed a distinct spatiotemporal sequence, peaking first in the cerebral cortex during early childhood (M: 6.6 [6.3–6.9] years, F: 5.4 [5.2–5.6] years, Δage = 1.2 years), followed by the cerebellum (M: 16.0 [15.3–17.1] years, F: 11.9 [11.2–12.7] years, Δage = 4.1 years) and subcortex (M: 16.8 [16.4–17.3] years, F: 14.7 [14.3–14.9] years, Δage = 2.1 years) during adolescence (all *P_perm_* _<_ 0.001, FDR-corrected; Fig. 2d–f). White matter volume (WMV) showed a more prolonged developmental trajectory, reaching its maximum in adulthood.

However, peak age did not significantly differ between sexes for cerebral WMV (M: 28.0 [27.2–28.6] years, F: 27.0 [26.1–27.7] years, Δage = 1 year; *P_perm_* _=_ 0.11; Fig. 2g) or cerebellar WMV (M: 27.1 [26.0–28.2] years, F: 24.6 [23.8–25.2] years, Δage = 2.5 years; *P_perm_* _=_ 0.02; uncorrected; Fig. 2h). Ventricular volume exhibited continuous nonlinear expansion across the lifespan, with no significant sex difference in peak age (Fig. 2i).

Global surface-based metrics revealed a pattern of sequential maturation: cortical thickness, the folding index, and surface area peaked in early, middle, and late childhood, respectively (Fig. 2j–l). The peak ages exhibited later patterns in males than in females for total surface area (M: 9.9 [9.0–10.6] years, F: 7.9 [7.2–8.6] years, Δage =2.0 years; *P_perm_* _<_ 0.001, FDR-corrected) but not for cortical thickness (M: 2.6 [1.5–3.1] years, F: 2.2 [1.7–2.5] years, Δage = 0.4 years; *P_perm_* _=_ 0.32) or the folding index (M: 5.8 [5.1–6.5] years, F: 5.8 [5.0–6.4] years; *P_perm_* _=_ 0.94).

In addition to sex differences in peak timing, males also exhibited larger absolute values than females across broad age ranges for most global measures (all *P_perm_* _<_ 0.005; Fig. S1). With respect to volumetric phenotypes (except for the total intracranial volume and ventricular volume), sex differences emerged in early childhood, reached their maximum magnitude during late adolescence to early adulthood, and gradually attenuated thereafter. With respect to surface-based phenotypes, sex differences in both surface area and the folding index emerged in early childhood but followed distinct trajectories: the differences in the folding index increased rapidly to peak at 6 years of age, whereas the differences in surface area continued to increase and reached their maximum in adulthood. Mean cortical thickness showed significant differences from approximately 15 years of age, reaching their maximum in adulthood and becoming not significant after 48 years.

### Sex-specific normative growth of subcortical regions

Subcortical regions followed inverted U-shaped trajectories in both sexes, reaching their peak volumes at different stages (Fig. 3, the left hemisphere). Specifically, the neostriatum showed the earliest peak maturation, with the caudate peaking in early childhood (M: 6.6 [6.0–7.1] years, F: 5.2 [4.8–5.6] years, Δage = 1.4 years; *P_perm_* _=_ 0.05, uncorrected; Fig. 3a), followed by the putamen in late childhood (M: 12.5 [11.9–12.9] years, F: 11.2 [10.7–11.7] years, Δage = 1.3 years; *P_perm_* _=_ 0.001, FDR-corrected; Fig. 3b). These were followed by the thalamus (M: 18.6 [18.0–19.1] years, F: 16.1 [15.6–16.7] years, Δage = 2.5 years), amygdala (M: 19.6 [19.0–20.1] years, F: 18.3 [17.7–18.8] years, Δ age = 1.3 years), hippocampus (M: 20.2 [19.6–20.7] years, F: 18.1 [17.6–18.6] years, Δage = 2.1 years), and pallidum (M: 21.4 [20.5–21.9] years, F: 19.2 [18.4–20.0] years, Δage = 2.2 years) (all *P_perm_* _≤_ 0.003, FDR-corrected; Fig. 3c–f), all of which reached peak volumes during adolescence or early adulthood (Fig. 3h). Absolute sex differences in these regions emerged primarily during the prenatal period or within the first two years of life (all *P_perm_* _<_ 0.005; Fig. S2), peaked in early adulthood and subsequently attenuated, although they remained detectable into late life (Fig. S2). Highly similar maturational patterns were observed in the right hemisphere (Fig. S3).

**Figure 3.**
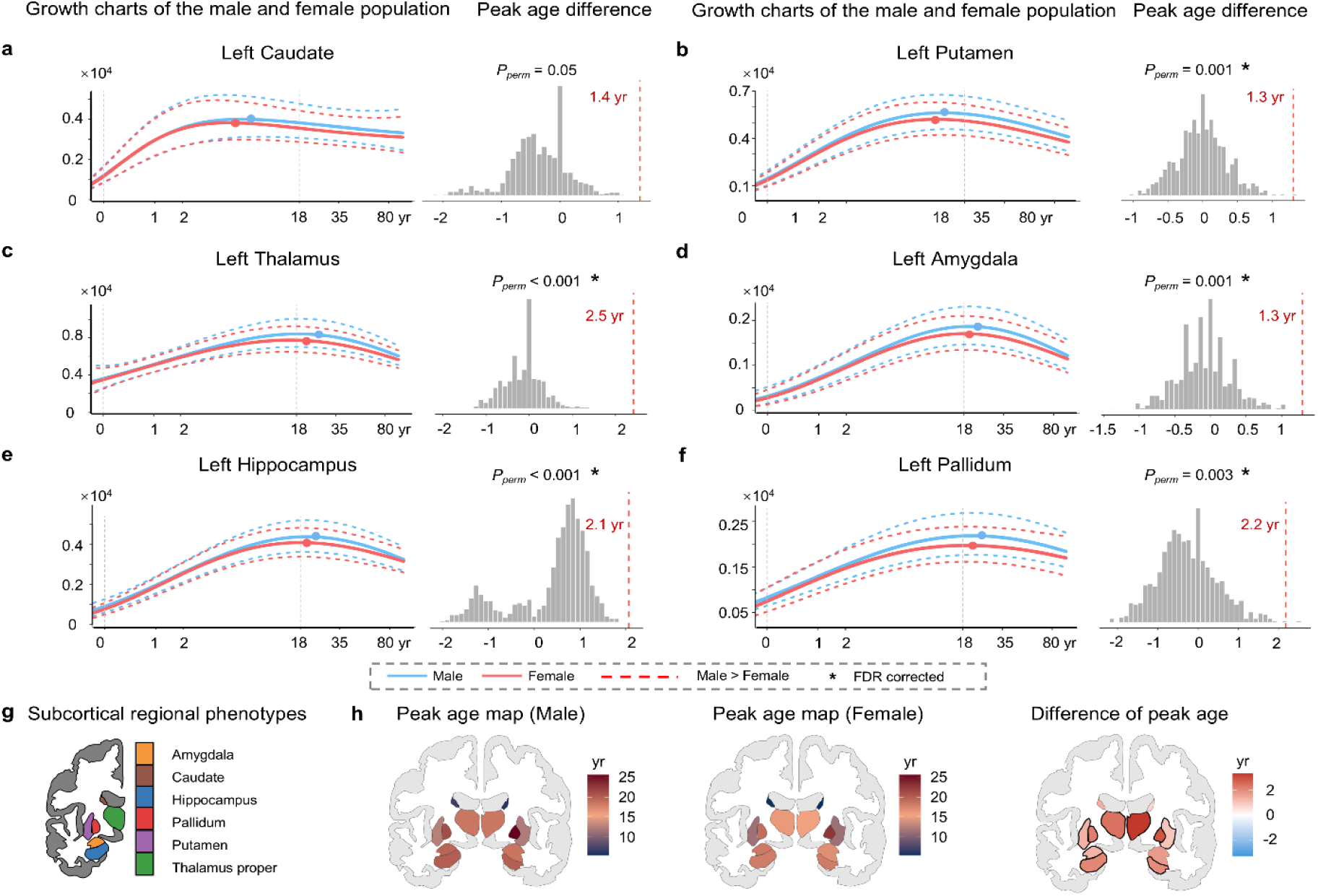
Sex-specific growth patterns of subcortical regional phenotypes. Growth curves for the subcortical volume of the caudate (**a**), putamen (**b**), thalamus (**c**), amygdala (**d**), hippocampus (**e**), and pallidum (**f**) in the left hemisphere. For each phenotype, the left side represents the normative growth curves for males and females (solid line: median; dotted lines: 5th and 95th percentiles), and the right side represents the null distribution of peak age differences evaluated via a two-sided permutation test (1,000 iterations; see Methods). **g**, Anatomical schematic of the subcortical regions in the left hemisphere. **h,** Peak age maps for male and female participants and the peak age difference map between sexes. The black contours indicate significant differences (*PFDR* < 0.05). The results for the right hemisphere are shown in Fig. S3. GMV, grey matter volume; yr, year.

### Sex-specific normative growth of cortical regions

We next mapped sex-specific lifespan trajectories of regional cortical phenotypes (Fig. 4a–c). The neurodevelopmental sequence was highly conserved between the sexes across these phenotypes. Mirroring global trends, regional cortical thickness peaked first, followed by the folding index and GMV, with the surface area peaking latest (Fig. 4a, b). Specifically, regional GMV showed the most widespread sex differences in peak timing, with males peaking 0.4–2.5 years later than females across 51 regions, spanning the medial frontal and parietal cortices, lateral temporal cortex, and insular cortex (all *P_FDR_* _<_ 0.05; Fig. 4c). The surface area also showed extensive male-specific delays, with a spatial topography similar to that of the GMV (all *P_FDR_* _<_ 0.05; Fig. 4c). With respect to cortical thickness, 13 regions with significant sex differences (including the prefrontal and medial parietal cortices) peaked later in males and were delayed by 0.6–2.0 years (all *P_FDR_* _<_ 0.05; Fig. 4c). In contrast, male delays in the folding index were restricted to a few regions in the lateral temporal and medial parietal cortices, although the right isthmus cingulate uniquely peaked earlier in males (all *P_FDR_* _<_ 0.05; Fig. 4c).

**Figure 4.**
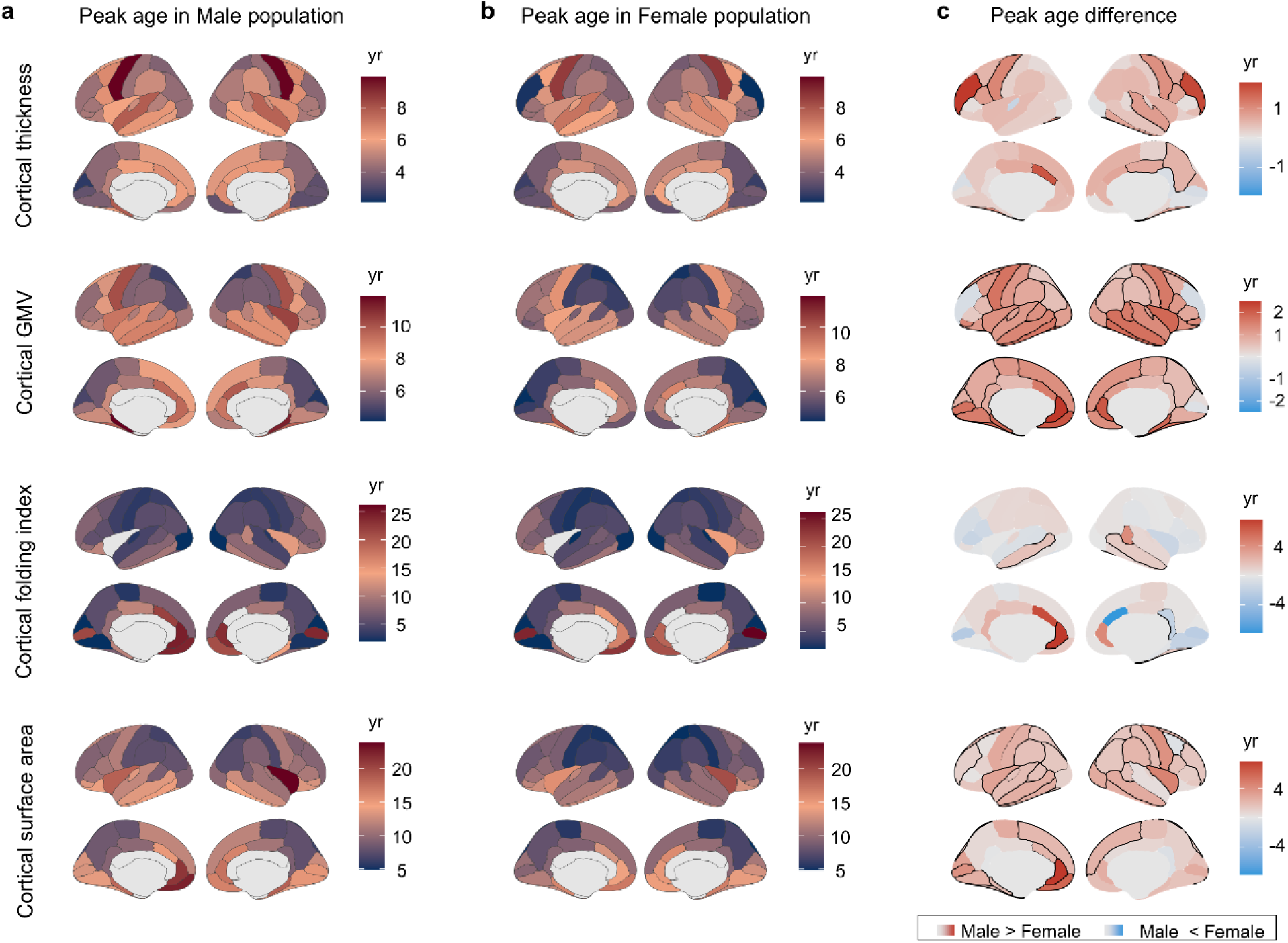
Sex-specific growth patterns of cortical regional phenotypes. Cortical maps showing peak age in males (**a**) and females (**b**) and the differences in peak age between males and females (**c**). From top to bottom, the four rows correspond to regional cortical thickness, cortical GMV, the cortical folding index, and cortical surface area, respectively. The black contours indicate significant differences in cortical regions (*PFDR* < 0.05). GMV, grey matter volume; yr, year.

Second, the developmental timing of absolute sex differences was highly phenotype-specific (Fig. S4 and Fig. S5). Significant sex differences (*P_perm_* _<_ 0.005) in surface area and GMV generally emerged during infancy or early childhood, whereas those in cortical thickness and the folding index appeared substantially later (Fig. S4). The timing of maximal sex differences also varied across phenotypes, occurring predominantly during adolescence and early adulthood for GMV and cortical thickness, but showing a more gradual increase that often extended into later adulthood for surface area and the folding index (Fig. S5).

### Improved phenotypic predictive accuracy using sex-specific growth models

To demonstrate the necessity of sex-specific growth models, we compared the out-of-sample predictive performance of sex-specific and sex-pooled models. The full lifespan dataset (N = 59,915) was divided into training and testing sets, with healthy controls (HCs) from the clinical case‒control cohorts serving as the testing set (N_male =_ 2,062, N_female =_ 1,852, aged 5–96 years). The remaining 27,698 male and 28,303 female participants composed the training set to retrain the sex-specific models. Using these 56,001 healthy participants, we constructed two size-matched sex-pooled training datasets to establish corresponding baseline models (Fig. 5a; detailed in Methods). To apply the retrained models to the testing set while accounting for site effects, we estimated site-specific statistical offsets and derived site-adjusted trajectories via a maximum likelihood approach ^42^. Individualized predicted values and deviation scores were then calculated for the tested HCs against these site-adjusted trajectories.

**Figure 5.**
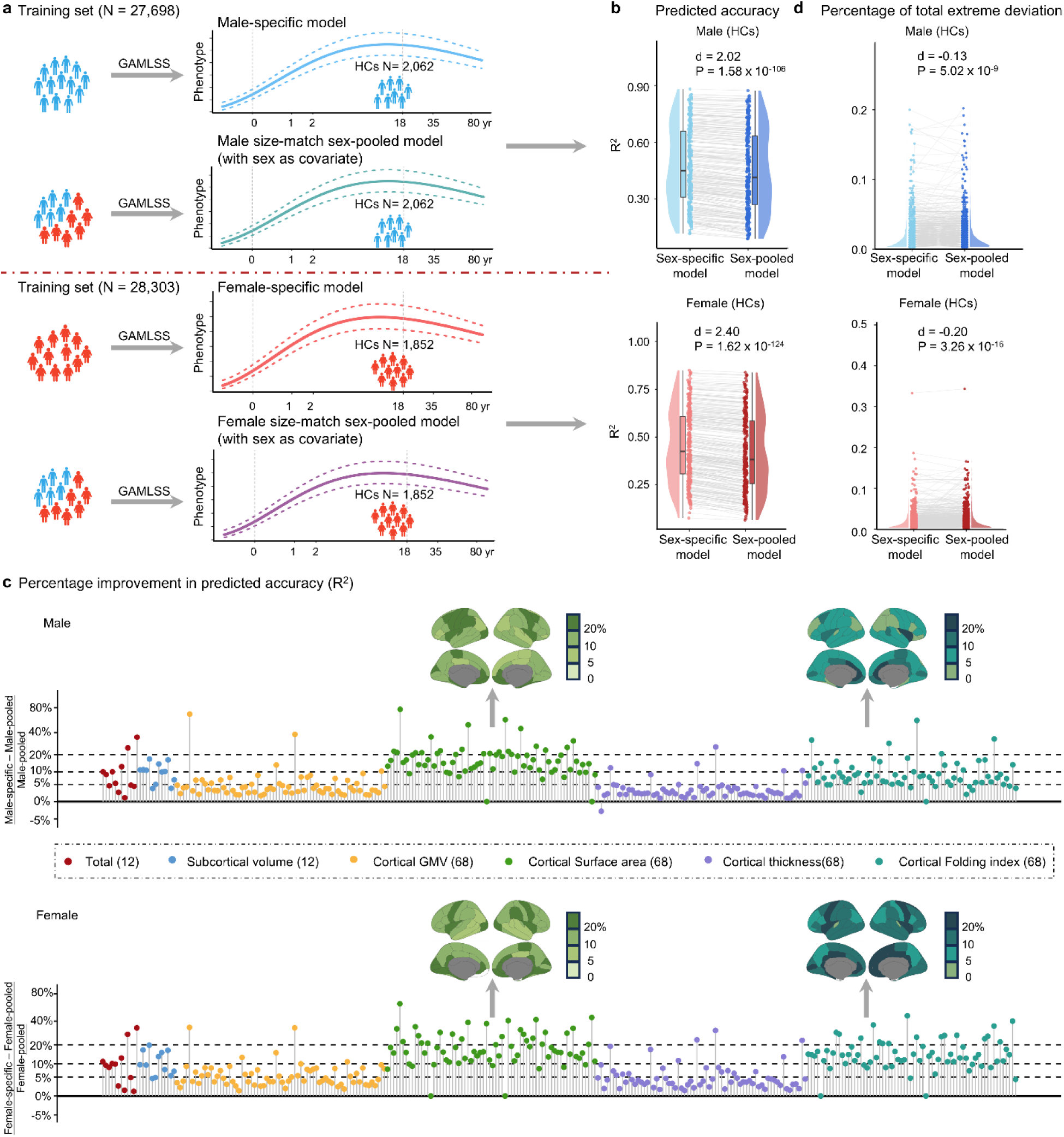
Out-of-sample predictive performance of sex-specific versus sex-pooled normative models in healthy individuals. **a**, Schematic workflow of the out-of-sample evaluation strategy. Healthy controls (HCs; 2,062 males and 1,852 females) from the clinical cohorts served as an independent testing set for individualized prediction. Sex-specific and size-matched sex-pooled models were retrained on the training dataset, comprising all healthy participants from the full lifespan dataset after excluding these held-out HCs. **b**, Predictive accuracy (R^2^) across the 296 structural brain phenotypes. Predictive performance is compared between sex-specific and size-matched sex-pooled models, evaluated separately in male and female held-out HCs. **c**, Relative improvement (%) in the predictive accuracy of sex-specific models over size-matched sex-pooled models. The cortical maps illustrate the spatial patterns of improvement in surface area and the folding index in both sexes. **d**, Differences in the percentage of extreme deviations in the testing set generated by sex-specific versus size-matched sex-pooled models. GAMLSS, generalized additive models for location, scale and shape; HCs, healthy controls.

We quantified the predictive accuracy (R^2^) for each phenotype (detailed in Methods), defined as the proportion of variance in observed phenotypes explained by the model in held-out HCs (Fig. 5b). The male-specific model outperformed the sex-pooled model (Cohen’s d = 2.02, *P* = 1.58×10^-106^), yielding accuracy improvements of > 10% for 96 phenotypes and > 5% for 163 phenotypes. The female-specific model demonstrated a similar advantage over the sex-pooled model (Cohen’s d = 2.40, *P* = 1.62×10^-1^^24^), with > 10% improvements in 134 phenotypes and > 5% in 197 phenotypes. The most substantial gains (> 20%) for these sex-specific models occurred in total WMV, alongside regional surface area and the folding index within the visual, cingulate, and insular cortices (Fig. 5c). Furthermore, we evaluated the out-of-sample calibration of the normative models by quantifying the proportion of total extreme deviations (|Z| > 2.6) in held-out HCs (detailed in Methods). Both the male-(Cohen’s d = -0.13, *P* = 5.02×10^-9^) and female-specific models (Cohen’s d = -0.20, *P* = 3.26×10^-16^) models yielded significantly lower proportions of extreme deviations than the sex-pooled model across the phenotypes did (Fig. 5d). This pattern held when positive and negative extreme deviations were analysed separately (Fig. S6). Together, these results indicate that sex-specific models reduce misestimation of brain deviations in healthy populations, thereby establishing a more accurate normative baseline for individualized assessment.

### Clinical relevance of sex-specific normative models

Next, we evaluated the clinical utility of the sex-specific normative models in five neuropsychiatric disorders, namely, AD (patients: N_male =_ 570, N_female =_ 529; HCs: N_male =_ 741, N_female =_ 897), MCI (patients: N_male =_ 1,705, N_female =_ 1,175; HCs: N_male =_ 735, N_female =_ 884), MDD (patients: N_male =_ 358, N_female =_ 504; HCs: N_male =_ 272, N_female =_ 388), ASD (patients: N_male =_ 612, N_female =_ 104; HCs: N_male =_ 517, N_female =_ 163), and ADHD (patients: N_male =_ 163, N_female =_ 57; HCs: N_male =_ 121, N_female =_ 128). We computed individual deviation *z* scores for 296 structural brain phenotypes using the abovementioned retrained sex-specific and sex-pooled models.

_G_iven the clinical importance of extreme deviations in characterizing neuroanatomical abnormalities ^42, 46–48^, we compared their detection rates between sex-specific and sex-pooled models (Fig. S7). With respect to AD, MCI, and MDD, the sex-specific models detected more positive extreme deviations in male patients (AD: Cohen’s d = 0.19, *P* = 9.12×10^-6^; MCI: Cohen’s d = 0.18, *P* = 2.57×10^-1^^3^; MDD: Cohen’s d = 0.33, *P* = 1.26×10^-9^) and more negative extreme deviations in female patients (AD: Cohen’s d = 0.30, *P* = 2.33×10^-11^; MCI: Cohen’s d = 0.28, *P* = 1.56×10^-21^; MDD: Cohen’s d = 0.32, *P* = 3.66×10^-12^). With respect to ASD, the sex-specific models revealed more positive (M: Cohen’s d = 0.29, *P* = 1.79×10^-12^; F: Cohen’s d = 0.22, *P* = 0.03) and negative (M: Cohen’s d = 0.17, *P* = 2.17×10^-5^; F: Cohen’s d = 0.24, *P* = 0.01) extreme deviations in patients across both sexes. In ADHD, these sex-specific models revealed more negative extreme deviations in male patients (Cohen’s d = 0.30, *P* = 0.0002), with no significant effect observed in female patients (Cohen’s d = 0.07, *P* = 0.61). Together, these findings highlight the necessity of sex-specific benchmarks for capturing sex-biased neuroanatomical patterns in patients with brain disorders.

Using these sex-specific normative models, for each disorder, we characterized the case–control differences in deviation scores separately for males and females and quantified whether disorder-related deviations differed between sexes. In AD patients, both female and male patients showed positive mean deviations (Fig. S8a) and case–control effects (Fig. 6a) in total ventricular volume and widespread negative mean deviations and case–control effects across most global (e.g., cortical/subcortical GMV and mean cortical thickness; Fig. 6a, Fig. S8a), subcortical (e.g., hippocampus; Fig. 6b, Fig. S8b), and cortical (e.g., lateral temporal and lateral/medial parietal cortices; Fig. 6c, Fig. S8c) phenotypes. Although these case‒control Cohen’s d maps revealed high spatial similarity between the sexes (r = 0.81–0.99, *P_spin_* _<_ 0.001, FDR-corrected; Fig. S9), female patients tended to exhibit significantly larger effect sizes across these phenotypes (*P*_FDR <_ 0.05; Fig. 6a–c), suggesting more severe structural atrophy than in males. We also observed distinct sex-divergent patterns: male patients showed positive case‒control effects on the folding index and surface area within the precentral gyrus and medial prefrontal cortex, whereas these effects were absent in female patients (Fig. 6c). Representative examples are shown (Fig. 6d), where disease-related deviations indicate female-biased atrophy (Cohen’s d_female <_ Cohen’s d_male <_ 0) in mean cortical thickness, left hippocampal GMV and left inferior parietal cortical thickness, and a male-biased increase (Cohen’s d_male >_ Cohen’s d_female >_ 0) in left precentral surface area. In MCI, patients exhibited a pattern similar to that of AD, although the effects were generally more modest (Figs. S10–11). MDD was characterized by predominantly negative case‒ control effects, with widespread cortical thickness alterations across prefrontal, temporal,_ p_arietal, and occipital cortices, whereas GMV alterations were more spatially restricted and primarily involved frontal and temporal cortices (Figs. S12–13). In contrast, ASD and ADHD showed significant disorder-related deviations in only a limited number of regions (Figs. S14–15). With respect to MDD, ASD, and ADHD, these deviations showed lower spatial similarity between sexes, and few sex-by-diagnosis interactions survived FDR correction (Fig. S9).

**Figure 6.**
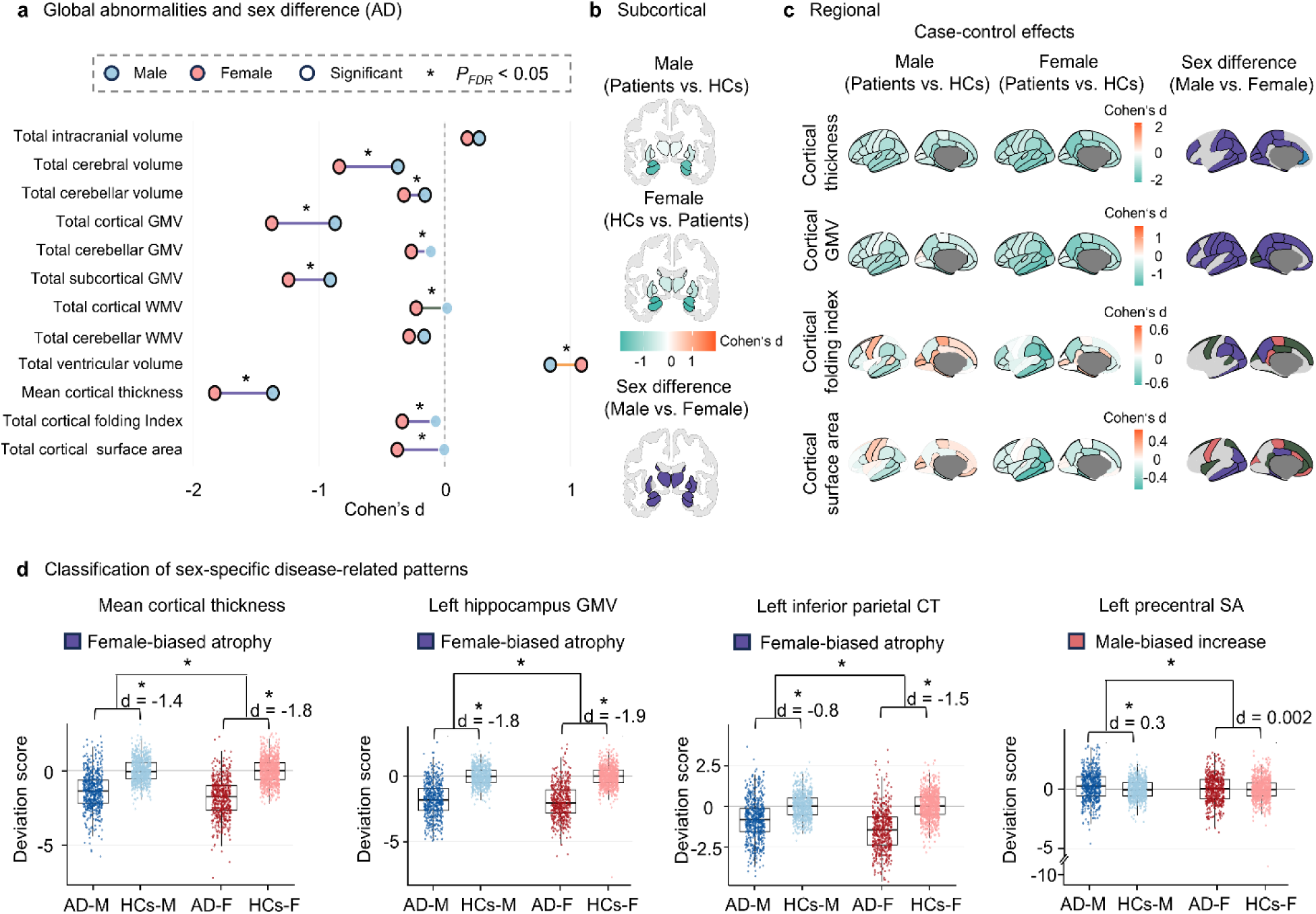
Sex-specific brain abnormality patterns in AD. **a**, Global structural abnormalities in patients. **b**, Subcortical structural abnormalities in patients. **c**, Regional cortical structural abnormalities in patients. In **a–c**, the columns show the male and female case–control effects (Cohen’s d) and categorical sex-difference patterns. In **a**, the blue and red circles represent the male and female case–control effects, respectively, and the black outlines indicate significant case–control differences (*PFDR* < 0.05). In **c**, the rows represent cortical thickness, GMV, the folding index, and surface area. The black contours indicate significant case‒control differences in cortical regions (*PFDR* < 0.05). Significant sex-difference patterns were classified according to the sex-by-diagnosis interaction and the corresponding male and female case‒control effects (see Methods). The corresponding results for the right hemisphere are shown in Fig. S16. **d**, Classification of sex-specific disease-related patterns. The boxplots show the distribution of deviation scores in patients and healthy controls for males and females. Sex-by-diagnosis interactions assess whether disease-related deviations differ between males and females. The asterisks (*) indicate *PFDR* < 0.05. Regions exhibiting significant sex-by-diagnosis interactions were classified according to the direction and relative magnitude of the male and female case–control effects (Cohen’s d), yielding distinct patterns of sex-specific disease effects. GMV, grey matter volume; CT, cortical thickness; SA, surface area; yr, year; HCs, healthy controls; AD, Alzheimer’s disease.

### _S_ensitivity analyses

To validate the robustness of the sex-specific lifespan growth trajectories and their peak age differences across all the phenotypes, we performed a suite of sensitivity analyses (for details, see the Methods section). Specifically, we evaluated (*i*) the stability of trajectories against sampling variations using bootstrap resampling (1,000 iterations), (*ii*) the potential bias of unequal sample sizes across age groups via age-stratified balance resampling (100 iterations), (iii) the influence of scanner sites using a leave-one-site-out (LOSO) approach (172 iterations), (iv) the reproducibility of the findings in independent subsets via a split-half strategy, and (*v*) the confounding effects of image quality by incorporating the Euler number as an additional covariate. Across all analyses, the resulting validation trajectories were highly consistent with the primary models for the vast majority of the phenotypes. To evaluate the robustness of the observed sex differences, we assessed the similarity between the peak age differences derived from each validation strategy and the main results. For strategies involving multiple iterations, we computed all possible pairwise peak age differences between the male and female validation models, with 1,000×1,000 cross-sex comparisons for the bootstrap analysis (*i*), 100×100 comparisons for the age-balanced resampling (*ii*), and 172×172 comparisons for the LOSO analysis (*iii*). For strategies (*iv*) and (*v*), the peak age differences were directly compared against the main findings. Finally, all sensitivity analyses reproduced the overall pattern of cross-sex peak age differences, yielding correlation coefficients of 1.00, 0.97, 0.99, 0.79, 0.76, and 0.80 for strategies (*ii*) through (*vii*), respectively (all *P* ≤ 2.90×10^-53^; Fig. S17).

## Discussion

Using large-scale neuroimaging data from 59,915 healthy participants spanning from 266 days postconception to 100 years of age, we constructed sex-specific growth charts for 296 structural brain phenotypes. These models revealed that males generally reached maturational peaks 1–2 years later than females did across most global and regional anatomical measures, with significant regional effects primarily concentrated in GMV and SA, whereas CT and FI exhibited relatively sparse and spatially restricted sex differences. This framework further provides a spatiotemporal map of when sex differences emerge and when they are most pronounced.

Compared with conventional sex-pooled models, the sex-specific models enhanced predictive accuracy, reduced the misestimation of individual deviations in held-out healthy participants and improved the detection of neuropathological deviations across five major neuropsychiatric conditions while capturing disease-related sex differences. Our work helps clarify sex-specific neurodevelopmental milestones and provides a tailored framework for sex-aware assessments of brain development, ageing and disease-related deviations.

Lifespan brain charts provide an important framework for understanding the normative principles of brain maturation across diverse neurobiological phenotypes, including structural measures ^42, 49–51^, cerebral perfusion ^52^, functional connectome ^43^, cortical similarity networks ^44^ and white matter micro-^53^ and macrostructure ^54^. Although recent landmark brain charts provide valuable insights, they typically treat sex as a fixed covariate ^42–44, 54, 55^, which inherently constrains male and female trajectories to share highly similar milestones. In addition, previous studies have reported sex differences in the timing of developmental peaks across various structural measures, but most of these studies were based on relatively limited sample sizes ^33–35^. More recently, a large-scale study ^56^ extended this framework by incorporating age-varying sex effects within a GAMLSS model, revealing significant sex biases across nearly all brain structures, converging with the results reported in our study. However, trajectory estimation remains influenced by data from both sexes through a shared model. In contrast, independently modelling male and female populations via stratified estimation removes this constraint. Here, we characterized sex-specific developmental trajectories and revealed robust sex differences in developmental milestones across the lifespan. Earlier maturation in females was observed across widespread brain regions, with the greatest sex differences concentrated in higher-order association areas. For instance, several regions involved in language processing, including the superior temporal gyrus, middle temporal cortex, and superior temporal sulcus, exhibited significantly earlier peak ages in females, with ages ranging from approximately 0.8 to 2.5 years. These findings are consistent with previous neuroimaging studies reporting earlier maturation of language networks in females ^57, 58^ and paralleling their earlier language acquisition and superior verbal fluency during childhood and adolescence ^59^. Earlier female peaks were also evident in regions involved in emotional processing, including the hippocampus, amygdala, insula and anterior cingulate cortex ^60, 61^. These differences may contribute to established sex differences in emotional reactivity and emotion regulation ^62, 63^. Beyond their functional relevance, these spatially patterned sex differences may also reflect regional variation in molecular architecture. For example, the peak ages of the fusiform gyrus across multiple morphometric measures were significantly earlier in females than in males (Fig. 4c). These findings align with those of a recent single-cell transcriptomic study showing highly region-specific sex differences in cortical gene expression and identifying the fusiform gyrus as one of the most sexually dimorphic regions ^64^. Together, these findings highlight the importance of delineating sex-specific developmental milestones for understanding the biological organization of human brain maturation.

Sex bias in biomedical modelling has increasingly been recognized as a pervasive problem, as biological standards, diagnostic thresholds, and predictive algorithms derived from male-dominated ^65^ or pooled cohorts are often applied indiscriminately to both sexes ^66^, potentially leading to the misestimation of disease risk or treatment efficacy in females ^65, 67, 68^. Neuroimaging-based normative modelling faces similar challenges. For instance, Elleaume et al. showed that sex imbalance in brain chart construction can significantly affect the estimation of individual deviation scores ^69^. Consistent with these observations, our results demonstrate that, compared with conventional sex-pooled models, sex-specific normative models reduced the misestimation of individual brain deviations in healthy participants and provided greater sensitivity to disease-related heterogeneity across five major neuropsychiatric conditions. For example, in the AD cohort, our sex-specific normative modelling revealed that compared with male patients, female patients showed more pronounced structural reductions, indicating greater vulnerability in females. These observations are consistent with previous clinical studies demonstrating that female patients exhibit more severe atrophy ^70, 71^, particularly in the frontal, parietal and temporal regions ^72, 73^, and experience more pronounced cognitive decline following disease onset ^74^. A similar female-biased spatial distribution of structural deviations was also observed in the aMCI group. These findings are further supported by established neuropathological and neuroimaging studies showing that AD pathology preferentially affects the transentorhinal and entorhinal cortices, hippocampus ^75, 76^, posterior cingulate cortex, precuneus, and other association cortical regions ^77, 78^. Notably, the regional abnormalities exhibiting female-biased reductions in our study were predominantly localized to these vulnerable regions, supporting the biological relevance of the greater female structural vulnerability observed in AD. These findings suggest that constructing sex-specific normative references is not merely a statistical refinement, but an essential step towards more accurate assessment of brain development and disease-related alterations ^45^.

Several limitations need further consideration. First, sex differences in brain development likely arise from the combined influence of genetic, hormonal, environmental and psychosocial factors. However, the relative contributions of these intertwined factors cannot be disentangled using the current data, which were limited to binary, self-reported biological sex. Second, although our dataset encompasses a broad age range from 266 days postconception to 100 years, sample sizes remained relatively sparse at the developmental extremes, particularly in the early postnatal period and among the oldest-old individuals aged over 90 years. Third, the cross-sectional nature of the data precludes direct tracking of individual developmental dynamics ^79^. Future densely sampled longitudinal datasets will be essential for more accurately mapping sex-specific nonlinear trajectories and maturational milestones.

## METHOD DETAILS

### Image quality control process

All structural MR images were evaluated using a unified four-stage quality control (QC) pipeline ^43^ that integrates automated algorithms with expert visual inspection. By systematically identifying and excluding scans containing structural artefacts, this standardized framework was implemented to ensure robust data consistency and reliability. Comprehensive details regarding these QC procedures can be found in our previous study ^43^ and at https://github.com/sunlianglong/BrainChart-FC-Lifespan/blob/main/QC/README.md.

### Structural MRI data processing

While a single, unified processing pipeline across all age groups would theoretically maximize methodological consistency, the dramatic anatomical variations across the human lifespan—especially during the perinatal and early infancy periods—render such uniformity impractical. Thus, consistent with our prior study ^43^, we adopted age-specific processing strategies for early developmental data while maintaining a harmonized approach for the remainder of the lifespan.

Specifically, for participants aged two years and older, all structural MRI data were processed via FreeSurfer v6.0.0, integrated within the containerized HCP structural preprocessing pipeline (v4.4.0-rc-MOD-e7a6af9) ^80^, deployed on the QuNex platform (v0.93.2) ^81^. Except the ABCD and UKB datasets, we applied the HCP-recommended PreFreeSurfer pipeline prior to FreeSurfer. This stage focused on normalization of anatomical images, which involved (1) brain extraction, (2) denoising, and (3) bias field correction of T1-weighted and, when available, T2-weighted scans. FreeSurfer processing was then conducted on the normalized images to generate cortical surfaces, which included (4) anatomical tissue segmentation, (5) construction of pial, white, and mid-thickness surfaces, (6) surface topological correction, and (7) projection of the surface onto a sphere for surface registration. For the structural images from the ABCD and UKB datasets, we used the officially released data and performed a processing procedure with FreeSurfer v6.0.0 ^82^.

For participants aged 0–2 years, we employed the iBEAT v2.0 pipeline ^83^. This tool is explicitly optimized for infant neuroimaging and has demonstrated enhanced accuracy in tissue segmentation and cortical reconstruction relative to alternative approaches ^83^. This pipeline follows the principles of the HCP-FreeSurfer approach and mainly includes (1) inhomogeneity correction of T1w/T2w images, (2) skull stripping, (3) cerebellum removal (for participants with incomplete cerebellum removal, frame-by-frame manual correction was performed), (4) tissue segmentation, (5) cortical surface reconstruction, (6) topological correction of the white matter surface, and (7) final reconstruction of the inner and outer cortical surfaces. For neonates from the dHCP study, we used data processed with the officially recommended dHCP pipeline ^84^, which was specifically designed to account for the substantial differences between neonatal and adult MRI data. This pipeline, which is similar in structure to those described above, mainly includes (1) bias correction and (2) brain extraction, which are performed on motion-corrected, reconstructed T2W images, (3) tissue segmentation, (4) cortical reconstruction of the white matter surface, (5) surface topology correction, (6) generation of pial and mid-thickness surfaces, and (7) spherical projection for surface registration.

Individual cortical surfaces for the 0–2 age group were mapped to the adult fsaverage standard space via a three-stage registration method ^43^. Briefly, for dHCP neonates (40 to 44 postmenstrual weeks), the following steps were implemented: (1) registering individual surfaces to their respective postmenstrual week templates ^85^; (2) aligning the 41–44-week templates to the 40-week baseline; and (3) registering the 40-week template to the fsaverage surface. Similarly, for infants aged 1–24 months, the following steps were taken: (1) registering the individual surfaces to their corresponding monthly age templates ^86^; (2) registering the monthly templates to the 12-month template; and (3) registering the 12-month template to the fsaverage surface template.

### Structural brain phenotypes

***(i) Global brain phenotypes***. Volumetric measures included total cerebral and cerebellar volume, ventricular volume and tissue-specific volumes for cortical/cerebellar GMV and WMV, as well as subcortical GMV. The surface-based metrics included the mean cortical thickness, total cortical surface area, and total folding index. Consistent with previous studies ^42^, total cerebral volume was defined as the aggregate of cortical GM and WM volumes—a proxy demonstrating a near-perfect correlation (r = 0.99) with direct segmentation. With respect to the 0–2-year age group, the iBEAT pipeline lacks precision in segmenting the cerebellum and ventricles; consequently, modelling for these specific phenotypes was restricted to participants aged 2–100 years.
***(ii) Subcortical regional phenotypes***. For participants aged 2–100 years, we utilized FreeSurfer segmentation to extract subcortical GM volumes, covering the bilateral thalamus, caudate, putamen, pallidum, hippocampus and amygdala. Conversely, for infants (0–2 years), volumetric measures were obtained by registering individual structural images to a 12-month template via age-appropriate registration techniques ^43^, followed by the mapping of a standard subcortical atlas.
***(iii) Cortical regional phenotypes***. Following surface reconstruction, we calculated vertex wise metrics including cortical thickness, GM volume, surface area, and the folding index for each participant. We utilized the individual-to-template surface mappings generated during preprocessing (see the “**Structural MRI data processing**” section) to project the standard Desikan–Killiany atlas onto each participant’s surface, thereby enabling the extraction of regionwise surface–based measures across 68 cortical regions. Previous research revealed that the entorhinal region, temporal pole region, and frontal pole regions exhibit excessive between-subject variability and outlier-level peak ages ^42^ because of signal artefacts in neonatal MR images. Consequently, these regions were excluded from the set of regional volumetric milestones in the lifespan brain chart proposed by Bethlehem et al. ^42^. To observe similar instability in our datasets, we excluded these three regions to ensure methodological consistency and result interpretability.

### Modelling the lifespan growth curves

We utilized generalized additive models for location, scale and shape (GAMLSS) ^87–89^—implemented via the *gamlss* package (version 5.4-3) in R 4.2.0—to estimate normative structural growth trajectories in the healthy population. Endorsed by the World Health Organization for normative modelling, GAMLSS constitutes a powerful statistical tool for delineating complex, nonlinear growth patterns, with broad applications in neurodevelopmental studies ^42, 43, 90^.

***(i) Sex-specific GAMLSS models***. Building on the framework established by Bethlehem et al.^42^, which identified the generalized gamma (GG) distribution as optimal for structural brain phenotypic data, we adopted this distribution characterized by the median (*μ*), coefficient of variation (*σ*), and skewness (*ν*). We determined the best-fitting parameters for each structural phenotype on the basis of this distribution. To flexibly model age-related changes, we employed fractional polynomial as smoothing term within the GAMLSS framework, allowing for the simultaneous fitting and selection of nonlinear terms. Optimization involved a systematic evaluation of model configurations, varying polynomial degrees (ranging from 1 to 3) for the μ and σ components of the GG distribution, and assessing the necessity of site-specific random effects. Consistent with Bethlehem et al. ^42^, we did not evaluate fractional polynomials for the *ν* parameter, as modelling *ν* in this manner frequently leads to instability (e.g., failure to converge to an optimal parameter) and lacks an a priori reason for assuming age- or random-effect-dependent changes in skewness. Model estimation utilized a default convergence threshold of log-likelihood = 0.001 between iterations, with a maximum of 400 iterations. We compared all the converged models using the Bayesian information criterion (BIC), selecting the configuration with the lowest BIC as the optimal model ^91^. Convergence was successfully achieved for all structural phenotypes. In the sex-specific framework male and female participants were modelled independently. Accordingly, sex was not included as a covariate, and separate normative models were estimated for each sex using identical model selection procedures. The final model specification for male total cerebral volume is provided below as a representative example.

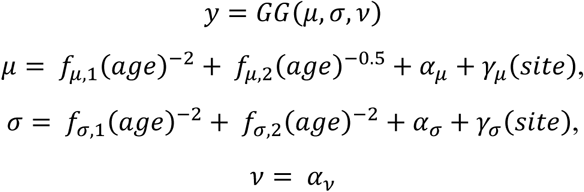

The *f* terms describe the fixed effects of age, modelled using fractional polynomials where the number of terms defines the polynomial order. In accordance with Bethlehem et al.^42^, age was quantified in gestational days and log-transformed to better capture the rapid structural changes during early development. Additionally, *α* represents the fixed intercepts, and *γ* represents site-level random effects. For the skewness parameter (*ν*), only a fixed intercept was included. Leveraging this GAMLSS framework, we generated male and female normative models for each phenotype on the basis of the sex-specific normative models.

***(ii) Sex-pooled GAMLSS models***. To provide a direct comparison with the sex-specific framework, we constructed pooled-sex normative models using the same GAMLSS pipeline, including the same GG distribution, fractional polynomial modelling strategy, parameter optimization procedure, BIC-based model selection, convergence criteria, and site-specific random effects. In contrast to the sex-specific framework, data from males and females were combined into a single training dataset, and sex was incorporated as a fixed-effect covariate in the *μ* and *σ* parameters. The final model specification for male size-matched (see the **“Construction of size-matched sex-pooled training datasets”** section) total cerebral volume is provided below as a representative example:

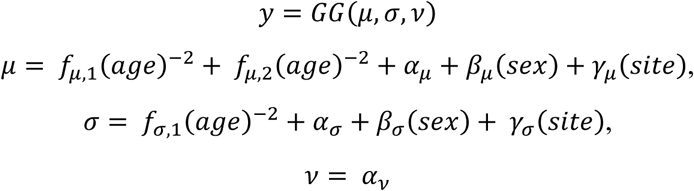

### Differences in peak age between the male and female growth curves

To evaluate whether the peak ages differed significantly between the male and female models, we employed a stratified permutation testing framework. First, the entire cohort of male (N=29,760) and female (N=30,155) participants was aggregated into a single dataset. This pooled sample was then segmented into 27 distinct age bins, ranging from prenatal stages to centenarians (-1–0 months, 0–3 months, 3–6 months, 6–12 months, 1–2 years, 2–4 years, 4–6 years, 6–8 years, 8–10 years, 10–15 years, 15–20 years, 20–25 years, 25–30 years, 30–35 years, 35–40 years, 40–45 years, 45–50 years, 50–55 years, 55–60 years, 60–65 years, 65–70 years, 70–75 years, 75–80 years, 80–85 years, 85–90 years, 90–95 years, and 95–100 years), to ensure granular age matching. Within each age stratum, sex labels were randomly shuffled while the original sample size ratios were maintained. This process effectively created two pseudogroups with randomized sex assignment but identical age distributions to the original data. For each permutation, normative growth curves were re-estimated for these pseudogroups, and the difference in peak ages was calculated. To construct a null distribution of peak age differences for each phenotype, this computationally intensive procedure was iterated 1,000 times. Two-tailed empirical p values were subsequently derived by locating the difference in observed peak age within this null distribution. Finally, to account for multiple testing, we applied a false discovery rate (FDR) correction at a threshold of 0.05, considering only those phenotypes that survived correction as exhibiting significant differences in peak timing.

### Sex differences in lifespan trajectories

To identify developmental periods during which male and female trajectories differed significantly, we performed pointwise permutation testing across the lifespan. The same stratified permutation procedure described above was used to generate a 1,000 pseudo-male and pseudo-female dataset by randomly shuffling sex labels within each of the 27 age strata while preserving the original age distribution and sample size of each sex. For every permutation, sex-specific normative trajectories were re-estimated using the same GAMLSS modelling pipeline. For each phenotype, the predicted population trajectories from the permuted male and female models were evaluated at 2^11^ uniformly sampled points along the transformed age axis, matching the resolution of the original normative curves. At each age point, the sex difference was calculated as the predicted male trajectory minus the predicted female trajectory, thereby generating a null distribution of trajectory differences across permutations. The observed true sex difference at each age point was then compared with this null distribution to obtain an empirical permutation p value. Developmental periods with *P_perm_* _<_ 0.005 were considered to exhibit significant sex differences in the predicted trajectories.

### Predictive accuracy for brain phenotypes of the male and female growth models

We evaluated the out-of-sample predictive performance of sex-specific models (comprising separate GAMLSS fits for males and females) against pooled models (a single GAMLSS fit with sex as a categorical covariate) using healthy controls from independent clinical sites. The validation dataset included 2,062 males (105 sites) and 1,852 females (98 sites) aged 5–96 years. Although these sites were part of the initial aggregate compilation, we rigorously removed them to create a leakage-free training set and retrained all the models on the remaining data. We then applied the retrained models to the held-out sites by estimating site-specific offsets using maximum likelihood, subsequently deriving individual centile scores for validation.

### Construction of size-matched sex-pooled training datasets

We constructed two size-matched sex-pooled training datasets that were matched to the male-specific and female-specific training cohorts. Matching was performed within each combination of study sites and predefined age intervals to ensure identical demographic compositions and age distributions between the sex-specific and sex-pooled models. Age stratification was performed using 27 lifespan intervals spanning prenatal stages to late adulthood: -1–0 years, 0–3 months, 3–6 months, 6–12 months, 1–2 years, 2–4 years, 4–6 years, 6–8 years, 8–10 years, 10–15 years, 15–20 years, 20–25 years, 25–30 years, 30–35 years, 35–40 years, 40–45 years, 45–50 years, 50–55 years, 55–60 years, 60–65 years, 65–70 years, 70–75 years, 75–80 years, 80–85 years, 85–90 years, 90–95 years, and 95–100 years. For each target sex-specific training cohort, we first calculated the number of participants within every unique StudyAgeGroup cohort, yielding a target sample size (Target_N)_ for each cohort. We then sampled participants from the pooled healthy population containing both males and females within the same StudyAgeGroup cohorts to construct the corresponding sex-pooled dataset. Sampling followed two predefined principles. First, the total number of selected participants within each StudyAgeGroup stratum was constrained to exactly match the corresponding Target_N f_rom the target sex-specific cohort. Second, within each stratum, participant selection was balanced towards an ideal 1:1 male-to-female ratio whenever possible. If one sex contained fewer available participants than needed, all available participants from that sex were included, and the remaining quota was filled using participants from the opposite sex. Using this procedure, we generated two independent size-matched sex-pooled datasets corresponding to the male-specific and female-specific training cohorts and subsequently trained two matched sex-pooled normative models for comparative analyses.

### Predictive accuracy of the normative model

The predictive accuracy of the normative models was evaluated using held-out healthy controls (HCs) from the clinical case‒control cohorts. For each phenotype, model-predicted values (*ŷ*) were compared against the observed phenotype values (*y*) using the coefficient of determination (*R^2^*). Crucially, the observed phenotype value (*y*) remained identical across analyses, whereas the predicted value (*ŷ*) differed according to the normative modelling framework. Specifically, predictions from the sex-specific framework were generated using independently trained male-specific and female-specific models, whereas predictions from the sex-pooled framework were generated using the corresponding size-matched sex-pooled models trained on pooled male and female participants. To ensure a fair comparison between modelling strategies, predictive performance was evaluated separately for males and females. For male HCs, the predictive accuracy of the male-specific model was compared against that of the sex-pooled model. Similarly, for female HCs, the predictive accuracy of the female-specific model was compared against that of the sex-pooled model. The predictive accuracy was quantified as *R^2^* = 1−SSE/SST, where SSE denotes the sum of the squared errors between the observed and predicted values and SST denotes the total sum of the squares relative to the mean of the observed values. To quantify the advantage of the sex-specific approach, we calculated the percentage improvement in predictive performance accuracy relative to the corresponding sex-pooled model. To avoid unstable estimates arising from near-zero baseline performance, this analysis was restricted to phenotypes with *R^2^* > 0.1 in the sex-specific and sex-pooled models. Improvement rates were computed as follows: 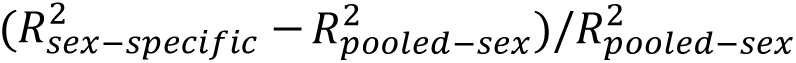. Finally, pairwise comparisons were performed to evaluate significant differences in accuracy between the models, reporting both effect sizes and significance levels.

Using the retrained normative models, all HCs were treated as full independent testing datasets for individual deviation estimation. For each participant, deviation scores (*Z* scores) were calculated relative to the corresponding sex-specific normative and size-matched sex-pooled models. Extreme deviations were defined as *z* > |2.6| (corresponding to *P* < 0.05). For each participant, we quantified the proportion of phenotypes exhibiting extreme positive, extreme negative deviations, and absolute extreme deviations.

### Clinical relevance of sex-specific normative models for brain disorders

To further evaluate the clinical applicability of the sex-specific normative models, we incorporated quality-controlled structural MRI data from six brain neuropsychiatric and neurodegenerative disorders. All preprocessing and quality control procedures were performed using an identical pipeline applied to the healthy dataset. The final clinical dataset included 1,099 patients with AD (male: 570 aged 53–92 years; female: 529 aged 49–95 years), 2,880 patients with MCI (male: 1,705 aged 42–93 years; female: 1,175 aged 48–93 years), 862 patients with MDD (male: 358 aged 12–65 years; female: 504 aged 11–77 years), 716 patients with ASD (male: 612 aged 5–62 years; female: 104 aged 5–54 years), and 220 patients with ADHD (male: 163 aged 7–21 years; female: 57 aged 7–20 years). Using the retrained normative models described in the “**Predictive accuracy for brain phenotypes of the Male and Female growth models**” section, all patient cohorts were treated as full independent testing datasets for individual deviation estimation. For each participant, deviation scores (*Z* scores) were calculated relative to the corresponding sex-specific normative trajectory after site adjustment. Extreme deviations were defined as *z* > |2.6| (corresponding to *P* < 0.05). For each patient, we quantified the proportion of phenotypes exhibiting extreme positive and extreme negative deviations and absolute extreme deviations.

### Regional pathological patterns and sex differences

To characterize sex-specific patterns of regional brain abnormalities, we compared deviation scores between patients and healthy controls (HCs) separately in males and females for each disorder. Analyses were performed independently for AD, MCI, MDD, ASD, and ADHD across CT, GMV, FI, and SA. For each brain region, two-sample independent t tests were conducted to compare deviation scores between patients and sex-matched HCs. Statistical significance was assessed separately for males and females. To quantify the magnitude of disease-related abnormalities, Cohen’s effect sizes were calculated for each region as follows:

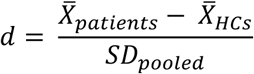

where *X̅_patients_* and *X̅_HCs_*_ _denote the mean deviation scores of patients and healthy controls, respectively, and *SD_pooled_*_ _represents the pooled standard deviation. Negative effect sizes indicate lower deviation scores in patients relative to controls, whereas positive values indicate higher deviation scores.

To evaluate the spatial similarity of these case–control abnormalities between the sexes, spatial correlation analyses were performed on the sex-specific case–control effects maps. For each disorder and across all six categories of brain morphometric metrics, Pearson correlation coefficients (r) were computed between the male and female case–control effects maps across the corresponding matched brain regions. The statistical significance of these spatial correlations was assessed using distinct frameworks depending on the type of features. For global and subcortical features, p-values were adjusted using the False Discovery Rate (FDR) method. For the four regional cortical metrics (cortical thickness, GMV, folding index, and surface area), which are subject to spatial autocorrelation, the significance of the spatial correlations was determined using spatial permutation tests (spin tests) ^92^.

To quantify sex differences in disease-related abnormalities, we constructed a two-way interaction model at the individual level using ordinary least squares (OLS) linear regression for each region of interest (ROI) as follows:

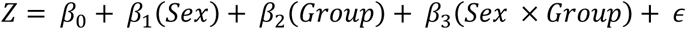

where Z represents the sex-specific, GAMLSS-derived normative deviation (Z score), Sex is dummy-coded with females as the reference group (female = 0, male = 1), Group is dummy-coded with healthy controls (HCs) as the reference group, and *ε* represents the random error term.

To elucidate the physical and statistical meaning of the interaction coefficient (*β*_3_), we examined the conditional expectations (predicted group means) of the *Z*-scores for the four subgroups as follows:

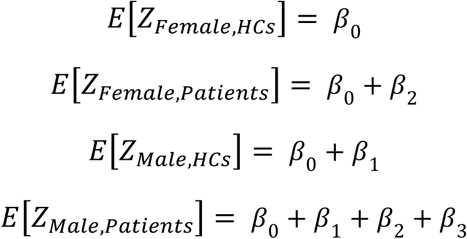

By calculating the disease-related alterations (patient minus control) for each sex, we obtain the following equation:

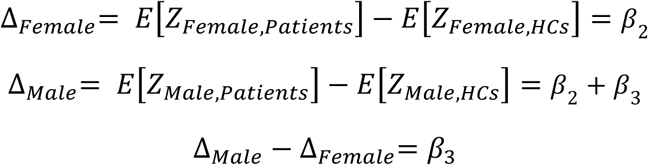

This algebraic formulation mathematically demonstrates that the interaction coefficient *β*_3_ purely quantifies the difference in disease-related deviation from sex-specific norms between females and males, with baseline normal sex differences (*β*_1_) fully controlled and cancelled out. The statistical significance of the interaction term was assessed using two-tailed t tests. Multiple testing correction was performed using the false discovery rate (FDR) across the 68 cortical ROIs within each morphometric modality (*P_FDR_* _<_ 0.05).

To facilitate biological interpretation of sex differences in disease-related abnormalities, ROIs showing a significant sex-by-diagnosis interaction (*P_FDR_* _<_ 0.05) were further classified into six categories by jointly considering the direction of the interaction coefficient (*β*_3_) and the signs of the sex-specific Cohen’s d values. Regions with negative Cohen’s d values in both sexes were classified as having a male-biased reduction (*β*_3_< 0) or a female-biased reduction (*β*_3_> 0). Regions with positive Cohen’s d values in both sexes were classified as having a male-biased increase (*β*_3_> 0) or a female-biased increase (*β*_3_< 0). Regions showing opposite directions of disease effects between sexes were classified as male reduction/female increase (males: Cohen’s d < 0; females: Cohen’s d > 0) or male increase/female reduction (males: Cohen’s d > 0; females: Cohen’s d < 0). These six categories were used to visualize sex difference patterns across global, subcortical, and cortical phenotypes.

### Sensitivity analysis of normative the growth models

The male and female normative growth trajectories, as well as their peak age differences, were validated using multiple sensitivity analyses. These analyses addressed key methodological concerns, including model stability, validation using age-matched subsamples with equal sample sizes across the two populations, potential site-specific effects, replication in independent subsamples, and assessment of the influence of imaging quality. For the sex-specific growth trajectories, we quantitatively assessed the similarity between these validated growth patterns and the main results by sampling each growth curve at 1,000 points, and computing the Pearson’s correlation coefficients between the corresponding curves. In the bootstrap resampling (1,000 iterations) and age-matched resampling (1,000 iterations) analyses, a small number of models occasionally failed to converge because of limited data being resampled from certain sites. These instances were resolved by repeating the resampling until convergence was achieved.

***(i) Bootstrap resampling analysis.*** To assess the stability of the lifespan trajectories and estimate confidence intervals, we conducted independent bootstrap resampling separately for the male and female cohorts. Using sampling with replacement, we generated 1,000 bootstrap datasets for each sex ^42^. Normative growth models were subsequently evaluated for within-sex model stability by quantifying the consistency between these bootstrap-derived trajectories and the original model. Furthermore, to verify the robustness of sex differences, we computed pairwise peak age differences for every combination of male and female bootstrap curves (a total of 10^6^ comparisons) and examined the correlation between these permutation-based differences and the primary findings across all the phenotypes.
***(ii) Age-matched sample sizes across two populations.*** To rule out the possibility that unequal sample sizes across age groups might drive cross-population differences, we performed a sensitivity analysis using age-stratified balance resampling. The data were divided into 27 age intervals (e.g., -1–0 months, 0–3 months, 3–6 months, …, 95–100 years; see the **“Peak age difference between male and female growth curves”** section). Within each interval, we randomly downsampled the majority sex to match the sample size of the minority sex, ensuring an equal number of male and female participants. We generated 100 such resampled datasets, with the total number of participants strictly fixed at 28,157 per population in each iteration. Subsequently, sex-specific normative models were refitted for all the phenotypes using these 100 datasets. We evaluated the models in two ways: First, to confirm within-sex stability, we calculated the similarity between each resampled growth trajectory and the original model. Second, to verify the robustness of the observed sex differences, we cross-compared the peak ages from the 100 male and 100 female models (yielding 10,000 pairwise comparisons). The correlations between these new pairwise peak age differences and our primary findings were then calculated across all the phenotypes.
***(iii) Leave-one-site-out analysis.*** To ensure that our lifespan trajectories were not driven by biases from specific imaging sites, we performed leave-one-site-out (LOSO) analyses across all the phenotype- and sex-specific models. During each iteration, data from one single site were removed, and the remaining data were used to refit the GAMLSS to generate a new growth trajectory. This process yielded 172 independent LOSO models each for the male and female cohorts. We then evaluated these models in two steps. First, to confirm within-sex stability, we quantified the similarity between each LOSO-derived trajectory and the original model. Second, to validate the robustness of sex differences, we extracted peak age differences across all possible pairs of the 172 male and 172 female LOSO trajectories. Finally, these pairwise differences were correlated with our primary findings across all the examined phenotypes.
***(iv) Split-half replication analysis.*** To evaluate how well our models could be replicated in independent datasets, we employed a split-half strategy. Participants from each sex were randomly divided into two subgroups, with the division strictly stratified by age and site. This yielded two subsets, each containing approximately 50% of the total sample (male: N₁ = 15,095, N₂ = 14,665; female: N₁ = 15,287, N₂ = 14,868). Within each newly formed subgroup, we independently re-estimated the lifespan normative growth trajectories for both males and females. Finally, across all the phenotypes, we evaluated the consistency of both the growth curves and the cross-sex peak age differences through three sets of comparisons: comparing each of the two subgroups against the primary model and comparing the two subgroups directly against one another.
***(v) Imaging quality–controlled modelling.*** To determine whether our lifespan growth trajectories were confounded by variations in image quality, we incorporated the Euler number—an established proxy for cortical surface reconstruction quality ^93^—into our GAMLSS framework as an explicit covariate for every phenotype and population. We subsequently evaluated the robustness of our primary findings by comparing the growth curves and peak age differences derived from these quality-adjusted models against those of the baseline models.

## RESOURCE AVAILABILITY

### Lead contact

Requests for further information and resources should be directed to and will be fulfilled by the lead contact, Yong He.

### Materials availability

This study did not generate new unique reagents.

### Data and code availability

The MRI dataset listed in Supplementary Table 1 are partially publicly available at the Adolescent Brain Cognitive Development Study (https://nda.nih.gov/), the Autism Brain Imaging Data Exchange Initiative (https://fcon_1000.projects.nitrc.org/indi/abide/), the ADHD200 (https://fcon_1000.projects.nitrc.org/indi/adhd200/), the Alzheimer’s Disease Neuroimaging Initiative (https://adni.loni.usc.edu/), the Age_ility Project (https://www.nitrc.org/projects/age-ility), the Baby Connectome Project (https://nda.nih.gov/), the Brain Genomics Superstruct Project (https://doi.org/10.7910/DVN/25833), the Calgary Preschool MRI Dataset (https://osf.io/axz5r/), the Cambridge Centre for Ageing and Neuroscience Dataset (https://www.cam-can.org/index.php?content=dataset), the Cincinnati MR Imaging of Neurodevelopment (https://nda.nih.gov/edit_collection.html?id=2329); the Consortium for Reliability and Reproducibility (https://fcon_1000.projects.nitrc.org/indi/CoRR/html/), the Dallas Lifespan Brain Study (https://fcon_1000.projects.nitrc.org/indi/retro/dlbs.html), the Developing Human Connectome Project (http://www.developingconnectome.org/data-release/second-data-release/), the University of North Carolina Early Brain Development Study (https://www.med.unc.edu/psych/research/psychiatry-department-research-programs/early-brain-development-research/), the FBIRN Phase II Multi-site fMRI programs (http://www.nbirn.net/Resources/Downloads), the Healthy Brain Network (https://fcon_1000.projects.nitrc.org/indi/cmi_healthy_brain_network/MRI_EEG.html), the Human Connectome Project (https://www.humanconnectome.org), the Lifespan Human Connectome Project (https://nda.nih.gov/), the Nathan Kline Institute-Rockland Sample Dataset (https://fcon_1000.projects.nitrc.org/indi/pro/nki.html), the Neuroscience in Psychiatry Network Dataset (https://nspn.org.uk/), the Open Access Series of Imaging Studies (https://sites.wustl.edu/oasisbrains/), the Pediatric Imaging, Neurocognition, and Genetics (PING) Data Repository (http://pingstudy.ucsd.edu/), the Pixar Dataset (https://openfmri.org/dataset/ds000228/), the Philadelphia Neurodevelopmental Cohort ( https://www.med.upenn.edu/bbl/philadelphianeurodevelopmentalcohort.html), the Queensland Twin Adolescent Brain (https://openneuro.org/datasets/ds004146/versions/1.0.4), the Strategic Research Program for Brain Sciences MRI Dataset (https://bicr-resource.atr.jp/srpbsopen/), the OpenfMRI database ds000115 (https://openfmri.org/dataset/ds000115/), the SchizConnect Project (including the BrainGluSchi, the COBRE, the MCICShare, and the NMorphCH dataset, https://schizconnect.org/), the Southwest University Adult Lifespan Dataset (http://fcon_1000.projects.nitrc.org/indi/retro/sald.html), the Southwest University Longitudinal Imaging Multimodal Brain Data Repository (http://fcon_1000.projects.nitrc.org/indi/retro/southwestuni_qiu_index.html), and the UK Biobank Brain Imaging Dataset (https://www.ukbiobank.ac.uk/). The dhcpSym surface atlases in aged from 40 to 44 postmenstrual weeks is available at https://brain-development.org/brain-atlases/atlases-from-the-dhcp-project/cortical-surface-template/. The UNC 4D infant cortical surface atlases are available at https://bbm.web.unc.edu/tools/. The fs_LR_32k surface atlas is available at https://balsa.wustl.edu/. The normative models of are shared online via GitHub (https://github.com/zhuminhui/Sex-specific-brain-charts).

Code for analyses is available via GitHub at https://github.com/zhuminhui/Sex-specific-brain-charts. Software packages used in this manuscript include MRIQC v0.15.0 (https://github.com/nipreps/mriqc), QuNex v0.93.2 (https://gitlab.qunex.yale.edu/), HCP pipeline v4.4.0-rc-MOD-e7a6af9 (https://github.com/Washington-University/HCPpipelines/releases), dHCP structural pipeline v1 (https://github.com/BioMedIA/dhcp-structural-pipeline), iBEAT pipeline v2.0.0 (https://github.com/iBEAT-V2/iBEAT-V2.0-Docker), MSM v3.0 (https://github.com/ecr05/MSM_HOCR), FreeSurfer v6.0.0 (https://surfer.nmr.mgh.harvard.edu/), FSL v6.0.5 (https://fsl.fmrib.ox.ac.uk/fsl/fslwiki), MATLAB R2018b (https://www.mathworks.com/products/matlab.html), R v4.2.0 (https://www.r-project.org), GAMLSS package v5.4-3 (https://www.gamlss.com/),and ggplot2 package v3.4.2 (https://ggplot2.tidyverse.org/). The codes for the GAMSS model are adjusted from Bethlehem et al. ^42^ at GitHub (https://github.com/brainchart/Lifespan).

## ACKNOWLEDGMENTS

This work was supported by grants from the scientific and technological innovation 2030 - the major project of the Brain Science and Brain-Inspired Intelligence Technology (2021ZD0200500 to Q. Dong), the National Natural Science Foundation of China (82021004 and 82327807 to Y. He, T24B2012 to L. Sun, 824B2051 to X. Liang), the Major Project of the National Social Science Foundation of China (24&ZD252 to Z. Han). We are grateful to the Adolescent Brain Cognitive Development (ABCD) Study, the Autism Brain Imaging Data Exchange (ABIDE) Initiative, the ADHD200 Initiative, the Alzheimer’s Disease Neuroimaging Initiative (ADNI), the Age_ility Project, the Baby Connectome Project (BCP), the Brain Genomics Superstruct Project (BGSP), the Calgary Preschool MRI Dataset, the Cambridge Centre for Ageing and Neuroscience (Cam-CAN) Dataset, the Cincinnati MR Imaging of Neurodevelopment Dataset (C-MIND), the Consortium for Reliability and Reproducibility (CORR), the Dallas Lifespan Brain Study (DLBS), the developing Human Connectome Project (dHCP), the University of North Carolina Early Brain Development Study (EBDS), the FBIRN Phase II Multi-site fMRI programs, the Healthy Brain Network (HBN), the Human Connectome Project (HCP), the Lifespan Human Connectome Project (HCPA & HCPD), the Nathan Kline Institute-Rockland Sample (NKI-RS) Dataset, the Neuroscience in Psychiatry Network (NSPN) Dataset, the Open Access Series of Imaging Studies (OASIS), the OpenfMRI database ds000115, the Pixar Dataset, the Philadelphia Neurodevelopmental Cohort (PNC), the Queensland Twin Adolescent Brain (QTAB), the Southwest University Adult Lifespan Dataset (SALD), the SchizConnect Project, the Southwest University Longitudinal Imaging Multimodal (SLIM) Brain Data Repository, the UCLA dataset, the UK Biobank (UKB) Brain Imaging Dataset, the Disease Imaging Data Archiving: major depressive disorder (DIDA-MDD) Working Group, and the Multi-center Alzheimer Disease Imaging (MCADI) Consortium.

ABCD: data used in the preparation of this article were obtained from the Adolescent Brain Cognitive Development Study (https://abcdstudy.org), held in the NIMH Data Archive (NDA). This is a multisite, longitudinal study designed to recruit more than 10,000 children age 9-10 and follow them over 10 years into early adulthood. The ABCD Study® is supported by the National Institutes of Health and additional federal partners under award numbers U01DA041048, U01DA050989, U01DA051016, U01DA041022, U01DA051018, U01DA051037, U01DA050987, U01DA041174, U01DA041106, U01DA041117, U01DA041028, U01DA041134, U01DA050988, U01DA051039, U01DA041156, U01DA041025, U01DA041120, U01DA051038, U01DA041148, U01DA041093, U01DA041089, U24DA041123, U24DA041147. A full list of supporters is available at https://abcdstudy.org/federal-partners.html. A listing of participating sites and a complete listing of the study investigators can be found at https://abcdstudy.org/consortium_members/. ABCD consortium investigators designed and implemented the study and/or provided data but did not necessarily participate in the analysis or writing of this report. This manuscript reflects the views of the authors and may not reflect the opinions or views of the NIH or ABCD consortium investigators. The ABCD data repository grows and changes over time.

ABIDE I: primary support for the work by Adriana Di Martino was provided by the (NIMH K23MH087770) and the Leon Levy Foundation. Primary support for the work by Michael P. Milham and the INDI team was provided by gifts from Joseph P. Healy and the Stavros Niarchos Foundation to the Child Mind Institute, as well as by an NIMH award to MPM ( NIMH R03MH096321).

ABIDE II: primary support for the work by Adriana Di Martino and her team was provided by the National Institute of Mental Health (NIMH 5R21MH107045). Primary support for the work by Michael P. Milham and his team provided by the National Institute of Mental Health (NIMH 5R21MH107045); Nathan S. Kline Institute of Psychiatric Research). Additional Support was provided by gifts from Joseph P. Healey, Phyllis Green and Randolph Cowen to the Child Mind Institute.

ADNI: Data used in preparation of this article were obtained from the Alzheimer’s Disease Neuroimaging Initiative (ADNI) database (adni.loni.usc.edu). As such, the investigators within the ADNI contributed to the design and implementation of ADNI and/or provided data but did not participate in analysis or writing of this report. A complete listing of ADNI investigators can be found at: http://adni.loni.usc.edu/wp-content/uploads/how_to_apply/ADNI_Acknowledgement_List.pdf. Data collection and sharing for this project was funded by the Alzheimer’s Disease Neuroimaging Initiative (ADNI) (National Institutes of Health Grant U01 AG024904) and DOD ADNI (Department of Defense award number W81XWH-12-2-0012). ADNI is funded by the National Institute on Aging, the National Institute of Biomedical Imaging and Bioengineering, and through generous contributions from the following: AbbVie, Alzheimer’s Association; Alzheimer’s Drug Discovery Foundation; Araclon Biotech; BioClinica, Inc.; Biogen; Bristol-Myers Squibb Company; CereSpir, Inc.; Cogstate; Eisai Inc.; Elan Pharmaceuticals, Inc.; Eli Lilly and Company; EuroImmun; F. Hoffmann-La Roche Ltd and its affiliated company Genentech, Inc.; Fujirebio; GE Healthcare; IXICO Ltd.;Janssen Alzheimer Immunotherapy Research & Development, LLC.; Johnson & Johnson Pharmaceutical Research & Development LLC.; Lumosity; Lundbeck; Merck & Co., Inc.; Meso Scale Diagnostics, LLC.; NeuroRx Research; Neurotrack Technologies; Novartis Pharmaceuticals Corporation; Pfizer Inc.; Piramal Imaging; Servier; Takeda Pharmaceutical Company; and Transition Therapeutics. The Canadian Institutes of Health Research is providing funds to support ADNI clinical sites in Canada. Private sector contributions are facilitated by the Foundation for the National Institutes of Health (www.fnih.org). The grantee organization is the Northern California Institute for Research and Education, and the study is coordinated by the Alzheimer’s Therapeutic Research Institute at the University of Southern California. ADNI data are disseminated by the Laboratory for Neuro Imaging at the University of Southern California.

BCP: data used herein is supported by NIH grant (1U01MH110274) and the efforts of the UNC/UMN Baby Connectome Project Consortium.

dHCP: data were provided by the developing Human Connectome Project, KCL-Imperial-Oxford Consortium funded by the European Research Council under the European Union Seventh Framework Programme (FP/2007-2013) / ERC Grant Agreement no. [319456]. We are grateful to the families who generously supported this trial.

HCP: data were provided by the Human Connectome Project, WU-Minn Consortium (Principal Investigators: David Van Essen and Kamil Ugurbil; 1U54MH091657) funded by the 16 NIH Institutes and Centers which support the NIH Blueprint for Neuroscience Research; and by the Mc-Donnell Center for Systems Neuroscience at Washington University.

HCP Lifespan: data used in this publication was supported by the National Institute of Mental Health of the National Institutes of Health under Award Number U01MH109589 and by funds provided by the McDonnell Center for Systems Neuroscience at Washington University in St. Louis. The content is solely the responsibility of the authors and does not necessarily represent the official views of the National Institutes of Health.

NKI-RS: funding for key personnel provided in part by the New York State Office of Mental Health and Research Foundation for Mental Hygiene. Additional project support provided by the NKI Center for Advanced Brain Imaging (CABI), the Brain Research Foundation (Chicago, IL), the Stavros Niarchos Foundation, and NIH grant P50 MH086385-S1.

NSPN: the NSPN study was funded by a Wellcome Trust award to the University of Cambridge and the University College London.

UK Biobank: this research has been conducted using data from UK Biobank (www.ukbiobank.ac.uk). UK Biobank is generously supported by its founding funders the Wellcome Trust and UK Medical Research Council, as well as the Department of Health, Scottish Government, the Northwest Regional Development Agency, British Heart Foundation, and Cancer Research UK.

## AUTHOR CONTRIBUTIONS

M. Zhu, L. Sun, X. Liang, and Y. He conceived and designed the study. L. Sun, X. Liang, and Y. He supervised the project. M. Zhu, L. Sun, X. Liang, and Y. He designed the methodology. L. Sun processed the data. M. Zhu analyzed the data. M. Zhu and L. Sun developed visualizations. M. Xia provide suggestions for the results interpretation. Y. Bi, Q. Cao, R. Chen, Y. Chen, T. Chen, Y. Cheng, Z. Dai, Y. Deng, Y. Ding, Q. Dong, J.-H. Gao, Q. Gong, Y. Han, Z. Han, C.-C. Huang, H. Huang, R. Huang, R. Huo, L. Li, Q. Lin, B. Liu, C. Liu, H. Liu, N. Liu, Ying Liu, Yong Liu, J. Lu, L. Ma, W. Men, C. Pang, S. Qin, W. Qin, J. Qiu, S. Qiu, T. Si, S. Tan, Y. Tang, S. Tao, D. Wang, F. Wang, J. Wang, P. Wang, Q. Wang, X. Wang, Y. Wang, D. Wei, Y. Wu, P. Xie, X. Xu, L. Yang, C. Yu, H. Yuan, H. Zhang, X. Zhang, G. Zhao, K. Zhao, and Y. Zheng collected a subset of the data for this study. M. Zhu wrote the first draft of the manuscript, with critical revisions from L. Sun, X. Liang, and Y. He. All authors reviewed the manuscript and approved the final version.

## DECLARATION OF INTERESTS

The authors declare no competing interests.

## Notes

### Competing Interest Statement

The authors have declared no competing interest.

